# Retrieving fMRI data in real-time: difficulties and pitfalls

**DOI:** 10.1101/2022.06.27.497807

**Authors:** Michael Lührs, Benedikt A Poser, Tibor Auer, Rainer Goebel

**Affiliations:** Faculty of Psychology and Neuroscience, Department of Cognitive Neuroscience, Maastricht University, The Netherlands; Maastricht Brain Imaging Center, Maastricht, The Netherlands; Brain Innovation B.V., Research department, Maastricht, The Netherlands; School of Psychology, University of Surrey, United Kingdom

## Abstract

One of the significant challenges in real-time fMRI environments is to ensure that the functional images are exported in real-time. The prerequired ability to reconstruct these images immediately after the acquisition has already been resolved in 2004. Nowadays, more sophisticated sequences allow for higher resolution and faster repetition times and thereby challenging the ability to export this data in real-time. In this article, we tackle the potentially arising problem of sending the reconstructed data from the MRI to an external PC to perform the real-time fMRI analysis. We show that depending on the implementation of the data transfer, long delays can occur that can differ drastically in time and how often they occur. In addition, we propose a solution for SIEMENS MRI devices which was tested and applied already on multiple MRI devices including 3T and 7T machines on different vendor software versions. This new technique can be used as a blueprint that can be directly applied to other manufacturers. We also provide the source code of the described solution and show that the delay in the data transfer can be significantly reduced to a tolerable level using our proposed procedure. Finally, we integrate measurement options for the data transfer times to improve quality measures in real-time fMRI environments (e.g., clinical) that can implement the proposed solution. Efforts should be taken by the real-time community and MRI manufacturers to employ a standardized real-time export e.g., similar to the lab streaming layer which is used as a standard export method in EEG environments.

## INTRODUCTION

A lot of different studies have shown the possible applications of real-time functional magnetic resonance imaging (rt-fMRI) and facilitate the importance of this method (Ciarlo et al., 2022; Lührs et al., 2019; Pandria, Athanasiou, Konstantara, Karagianni, & Bamidis, 2020; Russo, Lührs, Salle, Esposito, & Goebel, 2020; Sorger et al., 2009; Tursic, Eck, Lührs, Linden, & Goebel, 2020; Weber, Ethofer, & Ehlis, 2020; Weiskopf, 2012). The main prerequisite for successful rt-fMRI is the ability to get access to the fMRI data in real-time. Real-time can be seen as the time between two consecutive recorded images or in short, the repetition time (TR). One should, however, also note that this time interval should include both data export and processing to make sure that the application keeps up with the acquisition. The beforehand mentioned experiments can be seen as separate classes of applications for rt-fMRI: Quality assurance (QA), rt-fMRI analysis, brain-computer interfaces (BCI) and neurofeedback (NF); which will be discussed separately to highlight out the importance of reliable data transfer in different scenarios.

### Quality assurance

In a quality assurance design (Heunis et al., 2019, 2020; Lührs & Goebel, 2019), rt-fMRI data is used to verify that the recorded data has sufficient quality for subsequent use. This use can be done in real-time and offline scenarios. For offline scenarios, the rt-fMRI quality assurance can for example show that an experimental run needs to be repeated due to too much movement of the participant, both leading to low sensitivity of the neural signal of interest. For instance, the lack of significant activation could be because the participant did not correctly understand and follow the instructions or due to other technical issues in the experimental setup (e.g., screen not fully visible, auditory stimuli not audible, loss of synchrony between the acquisition and the stimulus presentation). More details about potential applications of real-time quality control can be found in (Heunis et al., 2020). Most of these measures don’t directly require that the data used for the measure is available in a stable real-time fashion since the data is not directly used for further real-time applications. Still the severity of a slow or unreliable real-time export can cause missing information which would potentially have helped to acquire better fMRI data.

### rt-fMRI analysis

This design has the potential to get fMRI analysis results in real-time which could for example be used to stop an experiment earlier as soon as enough data for a respective task is available. Also, this design can be a pre-step for the BCI and QA applications. The importance of a reliable data transfer lies here in the processing times needed in general for real-time preprocessing and analysis methods which would benefit a lot from fast transfer time allowing for more preprocessing and analysis time.

#### BCI

The field of BCIs includes a variety of approaches that use the fMRI data in real-time to determine different measures that can be used to build a non-invasive interface to the subject’s brain. The potential applications can include assessing consciousness, communication, controlling dynamic experimental design, and controlling an external device (Sorger & Goebel, 2020). The speed of the data export is particularly crucial for communication and controlling because it strongly influences the feasibility of the application.

#### Neurofeedback

A special technique within the BCI field is Neurofeedback (NF) which, in short, presents the current brain activity (implicitly or explicitly) to the participant and thereby allows for modulation of the respective brain region or pattern. The main goal of a NF training is to induce neural plasticity changes, which can be maintained after the experiment. Different methods are used to visualize the feedback signal. In this article, we mainly focus on the time-dependent variables of these experiments and differentiate continuous and intermittent designs. In continuous design, the feedback is presented in real-time continuously giving most of the information directly to the participant whereas in an intermittent design the participant receives the feedback after performing the respective task for a certain amount of time without receiving real-time feedback. The continuous feedback thereby requires that the respective fMRI data is available in real-time to be preprocessed and analyzed to generate the feedback signal. For intermittent designs, more time is generally available, but the duration of the intermittent designs can also be rather short still requiring reliable feedback presentation times.

As shown above, an essential part of the rt-fMRI procedure in addition to the acquisition, preprocessing, and potential QA, analysis, and BCI application is the transfer of the data, which serves as a glue between each step. In particular, the data transfer of the fMRI images from the scanner to an external processing platform for further processing of the data is crucial to guarantee a real-time scenario. Since there are multiple MRI device manufacturers and no standard format does exist, best to our knowledge, a standardized approach for the real-time data export would improve the quality and reproducibility. This would address the missing standardization in rt-fMRI experiments and ensure reliable transfer which is one of the significant requirements for rt-fMRI experiments. Without a reliable, stable, and fast data transfer the fMRI images can’t be analyzed in real-time which would cause potential incremental delays or discrepancies in, for example, the stimuli presentation times. This could result in missing data to provide meaningful information for the brain-computer interface (BCI). Most of the current rt-fMRI studies do not give enough detail about the underlying technical aspects of how the data is accessed and analyzed in real-time (Thibault, MacPherson, Lifshitz, Roth, & Raz, 2018). The same is true for any quality measures of the real-time data export concerning incremental delays or jittering during the data transmission or reconstruction. In this article, we investigate the potential difficulties and pitfalls that can occur during the export of fMRI data in real-time and point out the importance to resolve potential problems by directly providing solutions to the used procedures. This is done using a standard rt-fMRI setup by comparing different export procedures (“indirect”, “direct”) used simultaneously.

## METHODS

To investigate the influence of data transfer times in real-time fMRI experiments 27 datasets were acquired using a 3T MRI (Siemens Prisma 3T, Siemens Healthineers, Erlangen, Germany) as well as five datasets using a 7T MRI (Siemens Magnetom 7T, Siemens Healthineers, Erlangen, Germany) including different TRs but always 100 volumes. Detailed information is available in the respective references (Lührs et al., 2015; Lührs, Poser, Auer, & Goebel, 2017). To measure the data transfer times, the start of the volume acquisition, as well as the receive time of the specific volume on an external computer were recorded and two different data transfer procedures were investigated simultaneously by using both transfer methods at the same time. The data was transferred to a real-time analysis computer running Turbo-BrainVoyager (TBV, Brain Innovation B.V., Maastricht, The Netherlands) to log the processing times or a self-build receiving script (Available in python and MATLAB (Heunis et al., 2019)). The first procedure used a standard “indirect” export of single mosaic DICOM files for each volume separately using the underlying server message block (SMB) network protocol. We call this method “indirect” for easier differentiation between the two methods. This is provided by the MRI manufacturer without further tools needed. The (SMB) protocol was used since it is the default protocol implemented in Microsoft Windows desktop environments. Other protocols like SAMBA or NFS would result in a similar outcome. The second approach was a newly developed method based on a “direct” TCP/IP-based connection between the MRI reconstruction computer and the receiving external real-time computer which allowed to send the volume data and specific header information for each volume. In this case, the data was exported directly using a custom real-time data export module (image calculation environment (ICE) functor) that is appended to the end of the image reconstruction chain. An overview of the described procedure is shown in figure 1.

**Figure 1:**
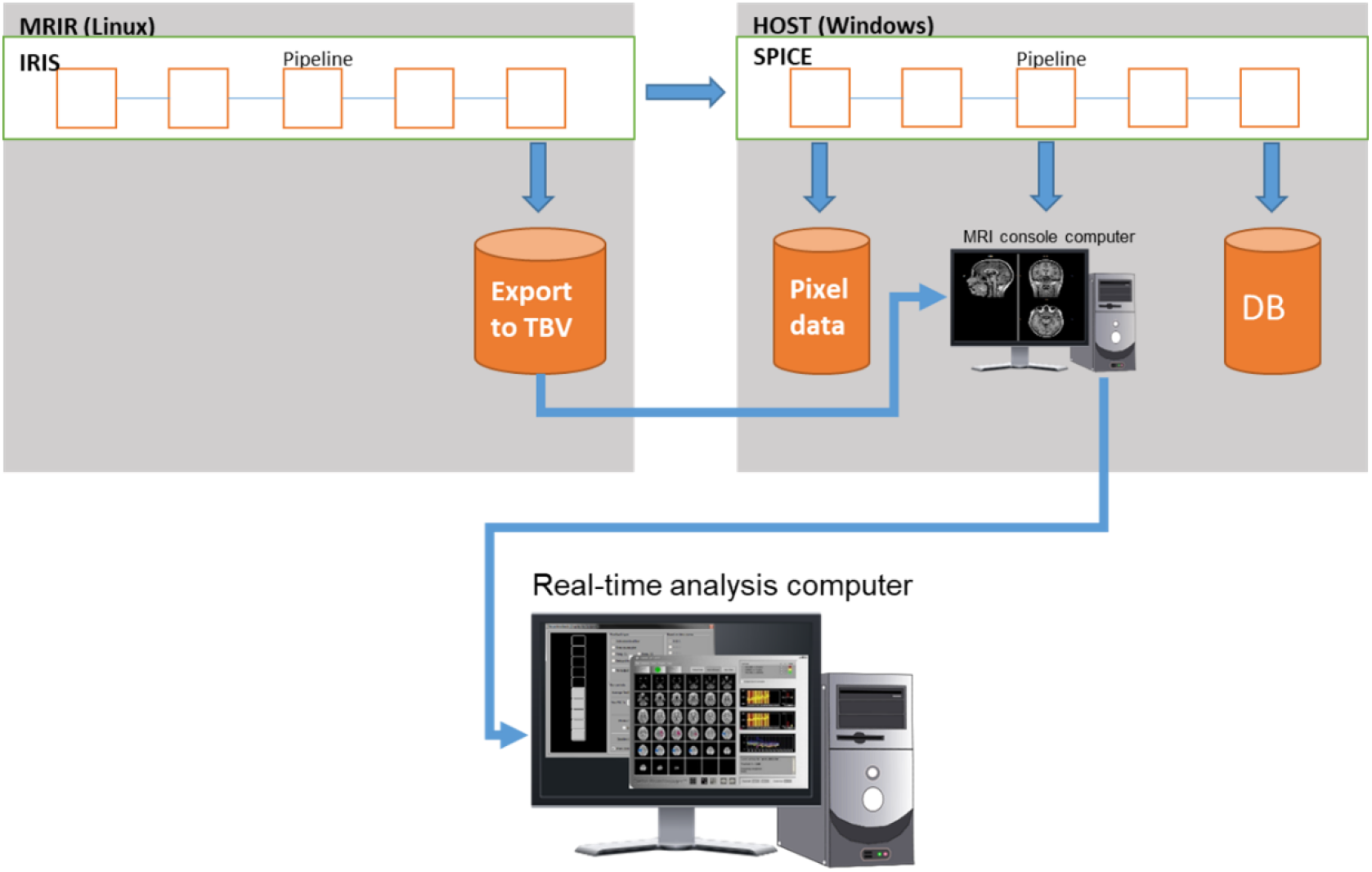
“Direct” real-time export procedure. In the “direct” real-time export procedure, the data is extracted using an ICE functor inserted as a last processing step in the reconstruction chain of the reconstruction server. The data is then directly transferred to the receiving real-time computer.

To connect the external real-time analysis computer to the image reconstruction computer a port forwarding tunnel was used which allowed to route the connection through the MRI host computer. The configuration of the port forwarding was performed using either the network shell (netsh (Microsoft, 2020)) from Microsoft Windows or a standalone port forwarding application (Putty) both running on the MRI host. In both procedures, the server that received either the DICOM or the pixel volume data was an external computer connected to the MRI host computer using a Gigabit-Ethernet connection. The measured times (volume acquisition trigger and volume receive times) were compared using the trigger at volume t+1 compared to receive time of volume t to calculate the pure data transfer time. Thereby the trigger was recorded separately to be most accurate.

## RESULTS

Overall, the mean data transfer time measured using the 3T MRI for the “indirect” file-based DICOM export was 513.9ms (+/-std 171.7ms) whereas the data transfer time for the “direct” TCP/IP-based export was 89.5ms (+/-std 76.9ms). For the 7T MRI, the “indirect” file-based DICOM export took 301.03ms (+/-std 87.14ms) compared to the “direct” TCP/IP-based export which needed 29.82ms (+/-std 18.29ms). The “indirect” file-based data export also showed strong jittering between successive volume data transfers which was not visible or less prominent in the TCP/IP-based export. The jittering was stronger for the multiband sequence measured on the 3T MRI at one point in time only regardless of the transfer method (vol 34). A general overview of the recorded data over time is shown in figure 2.

**Figure 2:**
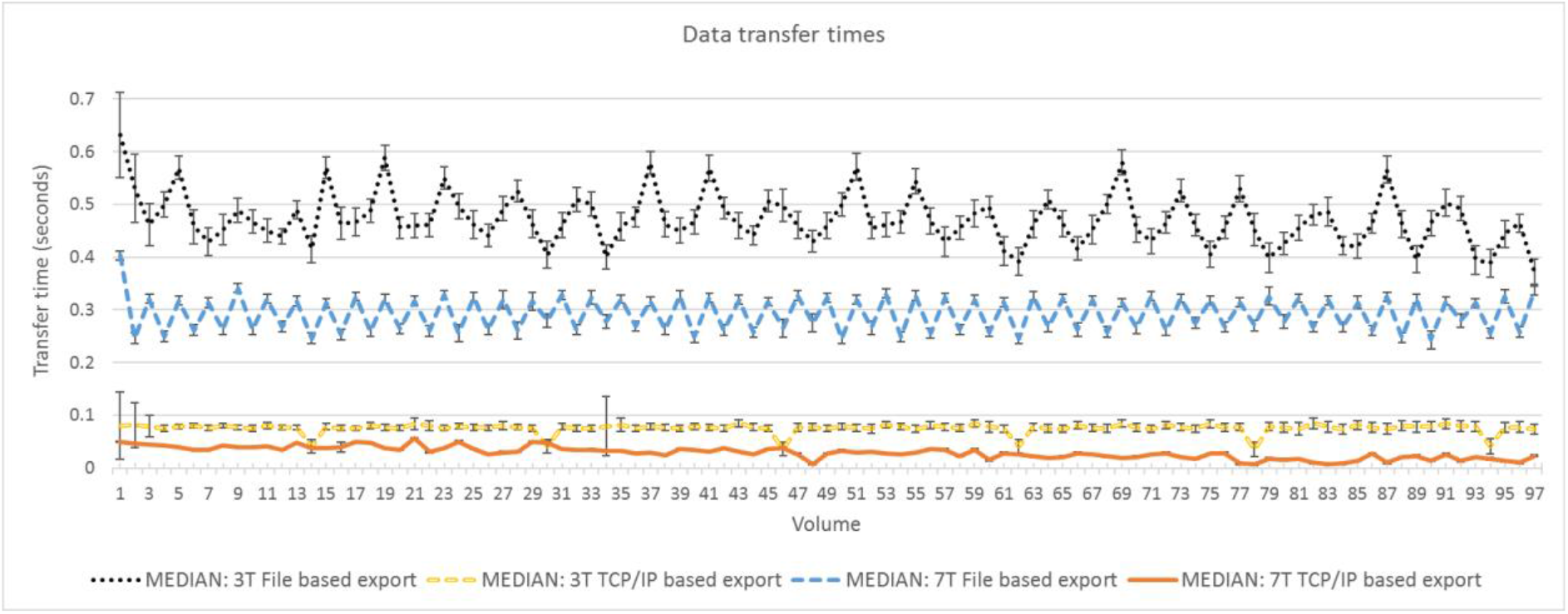
Median data transfer times using “indirect” files-based export and “direct” TCP/IP-based export on 3T and 7T MRI.

## DISCUSSION

The “direct” TCP/IP-based data transfer outperformed the “indirect” file-based export both in data transfer times as well as in the robustness of the data transfer, while the “direct” export leads to a decrease in transfer time variability. This can be seen in the much smaller averaged standard error as well as in the combined data transfer plots (see figure 2). Also, the jittering artifacts were more prominent in the file-based data export which could potentially be caused by the underlying network protocol SMB, which was designed to allow massive data exchange for many clients at the same time but seems to be not perfectly suitable for real-time export of fMRI data (especially earlier versions of the protocol). With newly developed sequences that allow acquiring high-resolution images (i.e. large number of voxels) while keeping the repetition times below 500ms, the file-based data export may not be able to keep up with the acquisition and an incrementally growing delay might occur which would violate the real-time condition in which the data must be retrieved and analyzed during one TR. In addition, such cutting-edge applications also require the more general elements of the system, such as network architecture, to be considered and optimized for the continuous flow of large amounts of data. Care must be taken when performing real-time fMRI experiments with ongoing feedback and fast sampling with very low TR not to overlook delays in the processing pipeline. Here we only investigated the data transfer time but not the reconstruction time itself, which could also be a potential problem for extremely low TR. Since we exported the data simultaneously using both methods, the reconstruction time would not influence our measures. It is reasonable to assume that use of both export methods in parallel did not by itself cause additional delays to either. Nevertheless, different sequences and acquisition protocols (especially spatial resolution and parallel imaging acceleration and multiband factors) result in different behaviors caused by the reconstruction itself. This was not systematically investigated in this work and should be further investigated.

### General guidelines

Simplifying the pipeline by avoiding unnecessary file write, transfer, and reading as implemented in a direct connection between the image reconstruction computer and the real-time analysis computer can improve quality assurance and help to ensure proper rt-fMRI data transfer and analysis. We only applied the solution using the MRI devices that were available to us, but the proposed solution can be generally applied to other manufacturers as well without further limitations. Since the proposed method might not be immediately available or suitable for all MRI devices (e.g. clinical scanners) we propose some guidelines below to potentially improve the “indirect” real-time data export.

#### The direction of file sharing

In the file-sharing procedure, one computer usually serves as a sender and the other one as a receiver. This also applies when sharing files through a shared folder using for example the SMB protocol. In this case, it is important to apply the correct order to prioritize file transfer. The computer that should perform the real-time computations on the fMRI data should create a shared folder on its system and the MRI console should connect to this shared folder and drop the fMRI images into the folder. This ensures that the file is listed immediately for transfer and acknowledged by the file transfer protocol.

#### SMB version

The version of the used network protocol is crucial since a lot of potential problems are resolved in later versions of the respective protocols. This is especially a problem for MRI devices since the protocols are part of the operating systems which are not always up to date for MRI console computers. If two computers are communicating the highest version available on both computers is used for file transfer. This could result in the use of an older version of the protocol, e.g. 2.0, which are slower and create more overhead per file transfer. To avoid any delay in name resolution, especially for older versions of the SMB protocol, the IP of the computer should be used to access the shared folder and write the file instead of the computer name.

#### Number of devices within the network

The more devices are in the local network the higher the chance that the file transfer might be delayed. Therefore, the network should in the best case only consist of the required computers. This could be achieved by using only the components in the network or creating a virtual network. Routing tables could also help to improve the transfer rates in the case of many devices in the same network.

#### Number of files to be transferred

Sending the whole functional data of one volume in one file is the optimal solution to reduce network delays. Splitting slices into separate files might reduce the performance and can cause additional delays. Still slice-by-slice transmission might be desirable depending on how the processing is performed (e.g. performing preprocessing on already available slices).

The developed “direct” approach does not need to consider the above-mentioned parameters because a direct TCP/IP connection is usually not affected by these problems. The only potential problem would be the bandwidth of the network interface. This should be able to handle the traffic to ensure a real-time transfer. The developed functor, as well as the source code, is available upon request.

### Quality assurance measures

To assure the data transfer in real-time is working correctly quality measures should be considered that point out the transfer time within each volume that is transferred. This could be implemented in many ways but would allow to interrupt the session and try to resolve the problem or to have the respective data transfer times available for post hoc evaluation. A standardized reporting of these transfer and processing times would be important for rt-fMRI experiments that use real-time data to present information to the participant. This is also true for intermittent designs since the delay of the data transfer can also reach several seconds (in the range of 10 to 20 seconds delay for a single volume).

### Potential consequences for rt-fMRI experiments

Since the problem is that the data transfer can be delayed in rt-fMRI experiments this can result in a variety of consequences. In case the transmission is jittered, and the data is not available regularly every TR can cause visual disturbances for the participant. Whether this influences the performance of the participant is unknown but should be investigated in more detail. For intermittent designs that need to display the data at a certain point in time, they could potentially fail to present the feedback since the data is not available. This could cause a general variability in the experiments and might be a problem in comparability across participants. Since the problem is not discussed widely in the community and not enough details are reported in the respective publications, the consequences are unknown but could potentially explain some cases for which for example the neurofeedback did not end up in a significant change in behavior. At least it is not possible to rule this out completely. As mentioned beforehand a reporting of these transfer and processing times would help to get more insights is the spread of the problem. In a variety of publications (Hampson et al., 2011; Hellrung et al., 2018; Krause et al., 2021; Marxen et al., 2016) a similar method for the direct data transfer are mentioned allowing to argue in both ways that the potential consequences are not as strong since solutions were already in place.

## CONCLUSION

We showed that standardizing the real-time export in rt-fMRI NF experiments is important to decrease the probability of errors caused by the variability in data transfer times. Additionally, we showed that the connection-oriented data transfer procedures of fMRI images lead to a decrease in transfer time variability which is especially important for NF experiments showing continuous feedback.

